# Dorsal raphe oxytocin receptors regulate the neurobehavioral consequences of social touch

**DOI:** 10.1101/850271

**Authors:** Z.A. Grieb, E.G. Ford, F.P. Manfredsson, J.S. Lonstein

## Abstract

Prosocial interactions are essential for group-living animals and are regulated by tactile cues shared among the group members. Neurobiological mechanisms through which social touch influences prosociality and related affective behaviors are relatively unknown. Using the evolutionarily ancient mother-young dyad as a model, we hypothesized that neurobehavioral consequences of social touch involves an interaction between central oxytocin (released during social touch) and serotonin (regulating affect and neuroplasticity). New mother rats showed upregulation of numerous aspects of the oxytocin system in the midbrain dorsal raphe (DR; source of forebrain serotonin) compared to non-maternal females. Preventing this upregulation by OTR knockdown in the maternal DR elicited infanticide, reduced nursing, increased aggression, and decreased active coping behavior. OTR knockdown also decreased serotonin-immunoreactive fibers, and increased neuroplasticity-restricting perineuronal nets, in the primary somatosensory cortex. Thus, oxytocin signaling in the DR regulates mechanisms involved in serotonin-induced cortical plasticity, which refines the tactile processing underlying prosocial behaviors.

---

Gentle touch is an essential, evolutionarily conserved determinant of social interaction and bonding in group-living animals^1–5^. It is so rewarding that adults of some highly social primate species spend up to 20% of their waking time prosocially touching each other (mostly grooming)^6^. The amount of time spent touching one another is even more striking between mothers and their young, with physical contact occurring for up to two-thirds of mothers’ time depending on the species^7–10^. The mother-offspring relationship is thought to have formed the basis from which other animal social relationships emerged^11–13^, so, the mother-offspring dyad provides an ideal model in which to study the neurobiological factors involved in how touch influences socioemotional behaviors. Neuropeptides, such as oxytocin, are known to be involved in prosocial touch and oxytocin is released centrally and peripherally when adults interact with each other or with infants^14–16^. Furthermore, oxytocin signaling is required for the numerous touch-associated behavioral modifications in new mothers, including their sensitive caregiving, protection of the young, and a positive affective state^17^.

Oxytocin may alter touch-associated maternal behaviors by acting on the midbrain dorsal raphe (DR), which is the source of most forebrain-projecting serotonin cells^18^. Experimental manipulations that enhance central serotonergic activity increase sociality and affect^19–21^, while manipulations that blunt serotonergic activity decrease sociality and affect^21–23^. Serotonin is also likely involved in the many neuroplastic changes occurring across pregnancy and lactation that are required for motherhood^24,25^. The DR has rarely been studied as a target of oxytocin’s effects on prosocial behaviors (or any other behaviors), despite the fact that the DR has long been known to express oxytocin receptors (OTRs)^26,27^. We propose that stimulating these OTRs during mother-offspring touch are essential for maternal socioemotional behaviors.

## Oxytocin is upregulated in new mothers

Female rodents become increasingly attracted to infants as pregnancy progresses, with a sharp peak around the time of parturition^28^. If OTRs in the DR are important for the expression of maternal socioemotional behaviors, then aspects of the oxytocin system in the DR, including OTRs and oxytocin fiber length, could be expected to show an upregulation across female reproduction; this would be similar to other brain regions involved in maternal behaviors^29–33^. In addition to the DR, the oxytocin system within the adjacent ventrolateral periaqueductal gray (PAGvl) may also show upregulation because the PAGvl, which contains the lateral wing subregion of the DR(DRlw)^18,34^, is sensitive to offspring sensory cues^35^ and densely innervates the DR^36^.

In support of our hypothesis, OTR autoradiographic binding in the rostral subregion of the DR was 2.5-fold higher in recently-parturient females compared to either diestrus virgins or postpartum day 7 mothers (Fig 1D). Female reproductive state did not significantly affect OTR binding in the other DR subregions studied or in the PAGvl/DRlw (Figs 1E-G). The higher OTR binding in the rostral DR of recently-parturient dams is likely due to the high circulating estrogens present at this time^37^, as there are positive relationships between estrogen receptor α and OTR expression in many other brain sites^38^. Estrogen receptor α in the DR is most expressed rostrally^39^, precisely where we found that female reproductive state increased OTR binding.

**Figure 1:**
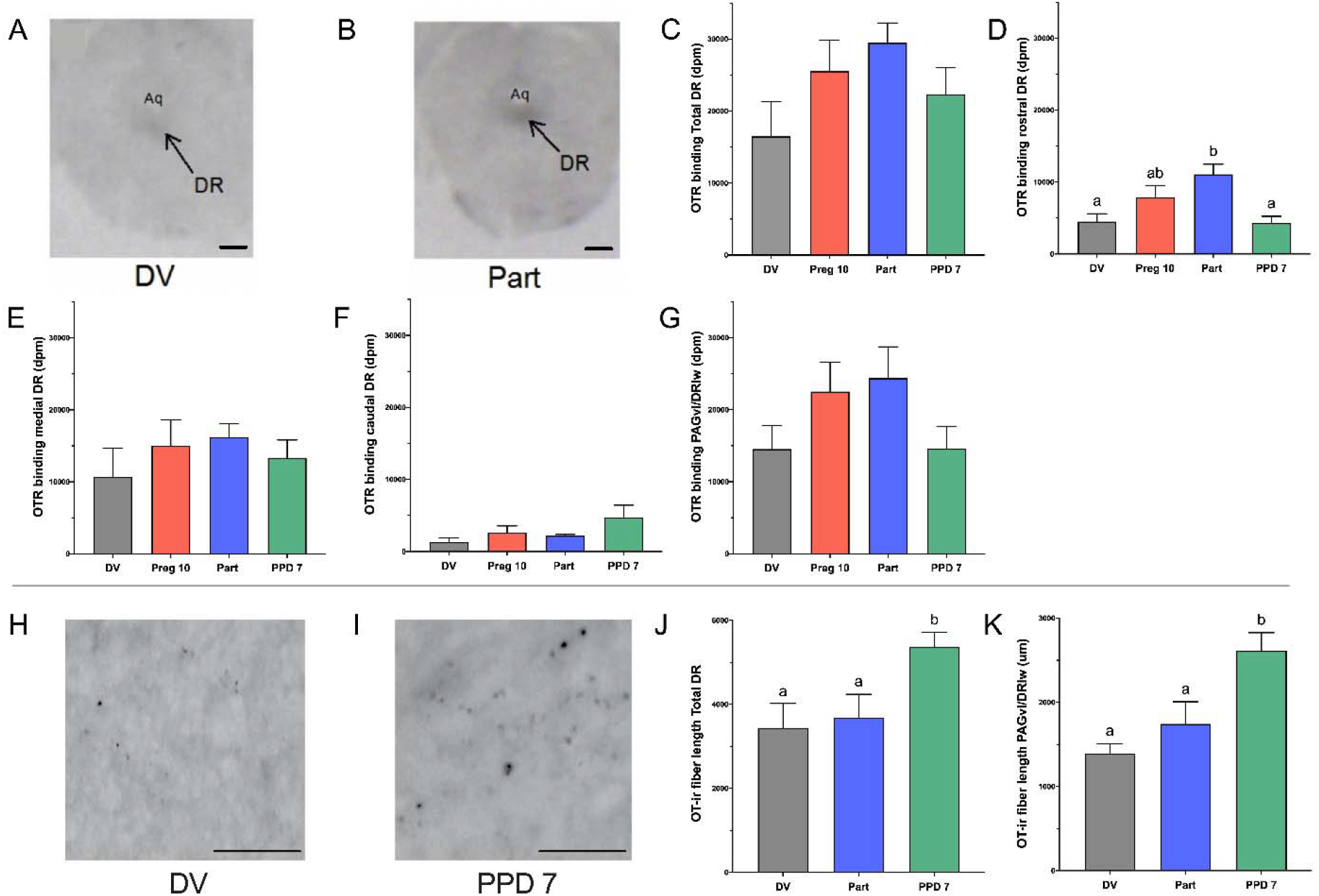
The oxytocin system is upregulated in new mothers. Representative autoradiographs of oxytocin receptor binding in the dorsal raphe (DR) of a (*A*) diestrus virgin (DV) and (*B*) recently-parturient (Part) rat. Oxytocin receptor binding in the (*C*) total DR, (*D*) rostral DR, (*E*) medial DR, (*F*) caudal DR, and (*G*) ventrolateral periaqueductal gray/DR lateral wing (PAGvl/DRlw) of females sacrificed as DV, on pregnancy day 10 (Preg 10), Part, or on postpartum day (PPD) 7. Representative photomicrographs of oxytocin-immunoreactive fibers in the DR of (*H*) DV and (*I*) Part females. Oxytocin-immunoreactive fiber length in the (*J*) DR and (*K*) PAGvl/DRlw of female rats sacrificed as DV, Preg 10, Part, or on PPD 7. Different letters above bars indicate significant differences between groups, *p* < 0.05. aq = cerebral aqueduct. Scale bars in A and B = 2 mm. Scale bars in H and I = 200 μm.

In addition to increased OTR binding, we also found greater total oxytocin-ir fiber length in the DR and PAGvl/DRlw of postpartum day 7 mothers when compared with either virgin or very recently-parturient females (Figs 1J, K). Oxytocin-ir fiber length reflects the capacity for local oxytocin release^32^, suggesting that mother rats with at least a week of caregiving experience have elevated capacity for oxytocin release in the DR and PAGvl/DRlw.

Given the higher level of OTR binding in the DR of recently parturient mothers, and the fact that estrogen receptor α is expressed on both serotonergic and non-serotonergic DR neurons^39^, we combined immunohistochemistry and *in situ* hybridization to determine the neuronal phenotypes in the DR expressing OTRs. Approximately 35% of the serotonergic neurons in the DR also expressed OTRs, although the number and percentage of serotonin cells expressing OTRs did not differ between recently parturient and virgin females (Figs 2E, F). Given that only about one third of the cells in the DR expressing OTRs are serotonergic^26^, we examined what other neuronal populations might also express OTRs. The second-largest population of neurons in the rat DR is GABAergic (at ~15%)^40^, which are mostly inhibitory interneurons. OTRs expressed on these cells may regulate tonic inhibition of serotonin released from the DR^41,42^. While recently-parturient mothers did not differ from virgins in the percentage of GABAergic neurons from all DR subregions that also expressed OTRs (Fig 2K), the percentage of GABAergic neurons in just the rostral DR that expressed OTRs was 44% lower in recently-parturient mothers than in virgins (Fig 2L).

**Figure 2:**
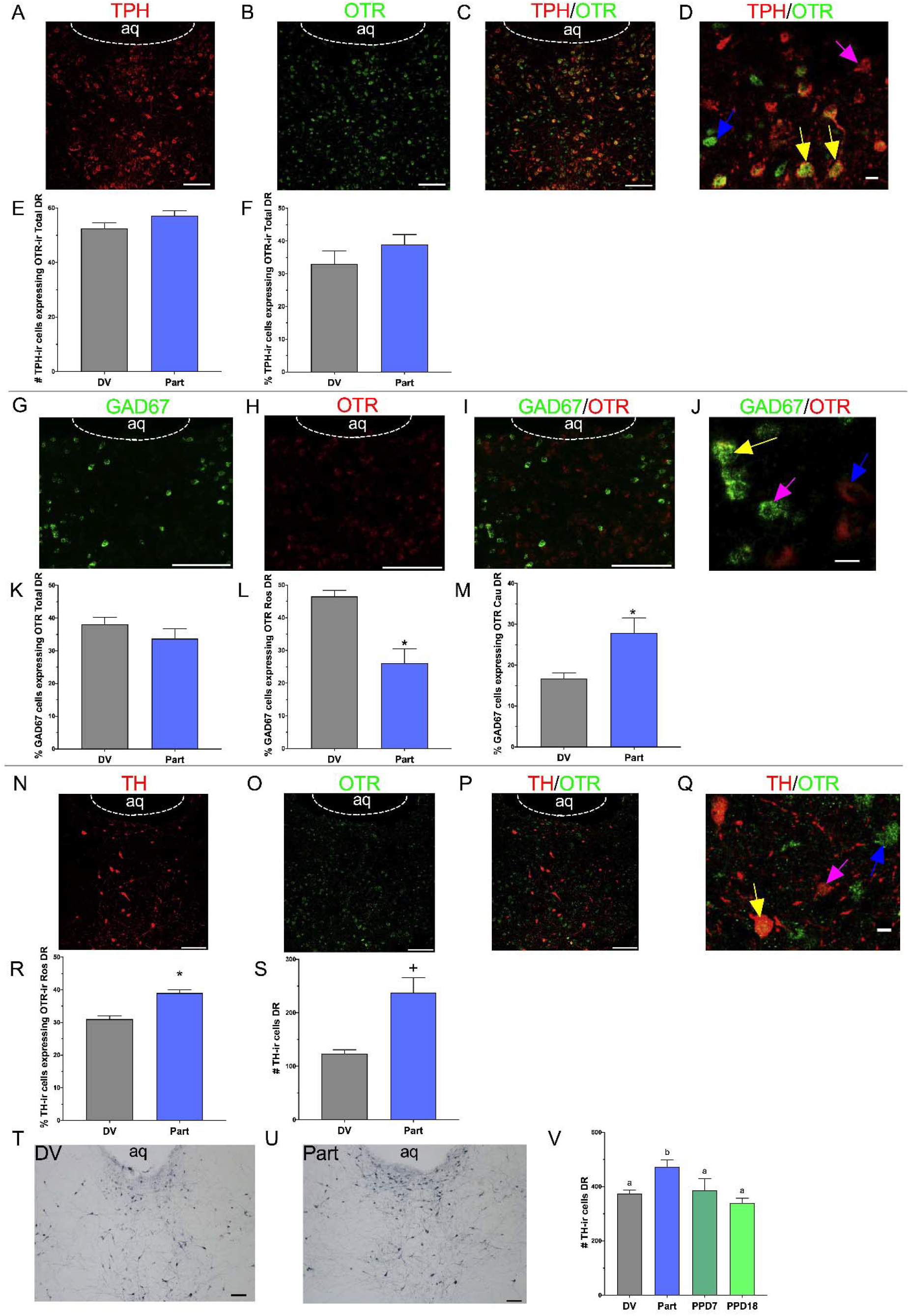
Oxytocin receptor expression is elevated in maternal midbrain serotonin and dopamine cells. Representative micrographs of (*A*) Tryptophan hydroxylase (TPH), (*H*) Glutamic acid decarboxylase^67^ (GAD^67^), (*N*) tyrosine hydroxylase (TH), (*B, H, O*) oxytocin receptor (OTR) immunofluorescence in the dorsal raphe (DR). TPH/OTR (*C, D*), GAD67/OTR (*I, J*) and TH/OTR (*P, Q*) overlapping immunofluorescence in the DR. Yellow, magenta, and blue arrows indicate colocalized, OTR-only, and TPH/GAD^67^/TH-only immunofluorescence, respectively. Number of (*E*) TPH/OTR and (*S, V*) TH cells in the DR of female rats sacrificed as diestrus virgins (DV), on pregnancy day 10 (Preg 10), as recently parturient dams (Part), or on postpartum day (PPD) 7. Percentage of (*F*) TPH cells in DR, (*K*) GAD^67^ cells in DR, (L) GAD^67^ cells in rostral DR, (*M*) GAD^67^ cells in caudal DR, and (*R*) TH cells in rostral DR also expressing OTRs in DV and Part females. * indicates *p* < 0.05. Different letters above bars indicate significant differences between groups, *p* < 0.05. aq = cerebral aqueduct. Scale bars in A-C, G-I, N-P, T, and U = 100 μm. Scale bars in D, J, and Q = 10 μm.

The DR also contains a fairly large, but often ignored, population of dopamine neurons (~1000 cells in rats)^43^. These cells have numerous similarities to the dopamine cells in the nearby ventral tegmental area, but, particularly relevant to the current studies, dopamine cells in the DR uniquely respond to social isolation^43^. This capacity to respond to social isolation could be essential for mothers’ motivation to be in contact with young rather than spend time alone. For both virgin females and recently-parturient mothers, we found that approximately one third of the cells across the DR that were immunoreactive for tyrosine hydroxylase (TH; dopaminergic marker) also expressed OTRs (Fig 2R). Both the number and percentage of TH-ir cells in the DR expressing OTRs were higher in recently-parturient mothers compared to diestrus virgins (Figs 2R, S). A follow-up analysis demonstrated that recently-parturient mothers also had more TH-ir neurons in the DR (regardless of OTR expression) compared to either diestrus virgins, postpartum day 7, or postpartum day 18 mothers (Fig 2V).

So far we have shown greater OTR binding in the rostral DR soon after parturition and greater oxytocin-ir fiber length shortly after, along with a greater percentage of rostral DR dopamine cells expressing OTRs (Figs 1D, 1J, 2R). These results collectively suggest that oxytocin has a greater influence on dopaminergic DR cells around the time of parturition, when maternal caregiving is first expressed. At the same time, recently-parturient mothers had a lower percentage of rostral DR GABAergic cells expressing OTRs (Figs 2L), suggesting that oxytocin has less influence on these inhibitory cells during the peripartum onset of maternal caregiving. The net result predicted from these findings is greater oxytocin-mediated stimulation, and less oxytocin-mediated inhibition, of serotonin and dopamine cell activity and probably release from the rostral DR of new mothers. This could presumably occur in response to either their naturally high endogenous oxytocin or after administration of exogenous oxytocin.

The serotonin neurons of the rostral DR preferentially project to many regions of the cerebral cortex^44,45^, suggesting that oxytocin’s effects on DR cells may selectively alter cortical serotonin release. Given that enhanced serotonergic activity increases socioemotional responses to touch^19–21^, increased serotoninergic output from the DR might be important for maternal socioemotional behaviors. On the other hand, many of the dopamine cells of the DR project to the same brain regions receiving innervation from the ventral tegmental area. Thus, their activation by oxytocin may increase maternal motivation and time spent with the young. Indeed, optogenetic activation of these DR dopamine neurons has been found to increase social investigation by inducing a negative affective state that can be alleviated by social interaction^43^. Therefore, oxytocin signaling in the rostral DR could regulate maternal socioemotional behaviors by: 1) increasing serotonin release into the cortex in ways that enhance sensitivity to social touch^19–22^ and 2) facilitating dopamine-mediated affective states that increase the motivation for prosocial contact^43^.

## Midbrain OTRs regulate maternal socioemotional behaviors

To investigate whether OTRs in the DR are *required* for maternal socioemotional behaviors, we injected an adeno-associated virus expressing a shRNA targeting OTR mRNA (Fig 3A) into the DR on pregnancy day 8, before any reproduction-related adaptations in female socioemotional behaviors occur^46,47^. When tested *in vitro*, our shRNA vector produced an almost 80% knockdown in OTR mRNA whereas the scrambled vector had no effect (Supplementary Fig 1A). When tested *in vivo*, injecting the shRNA vector into the DR on pregnancy day 8 resulted in approximately 60% knockdown in OTR mRNA when animals were sacrificed within 3 hrs after parturition (Fig 3C). Importantly, our OTR knockdown vector did not affect vasopressin V1a receptor expression (Supplementary Fig 1B) or the total number of serotonergic cells in the DR (Supplementary Fig 1D). After parturition, we observed a collection of social and affective behaviors including mothers’ undisturbed caregiving behaviors each day, maternal motivation using pup retrieval tests, anxiety-like behavior in an elevated plus maze and light-dark box, postpartum aggression in a resident-intruder paradigm, and anhedonic and active coping behaviors using saccharin preference and forced swim tests, respectively (Fig 3A). We predicted that knocking down OTRs in the DR would interfere with most of these behaviors.

**Figure 3:**
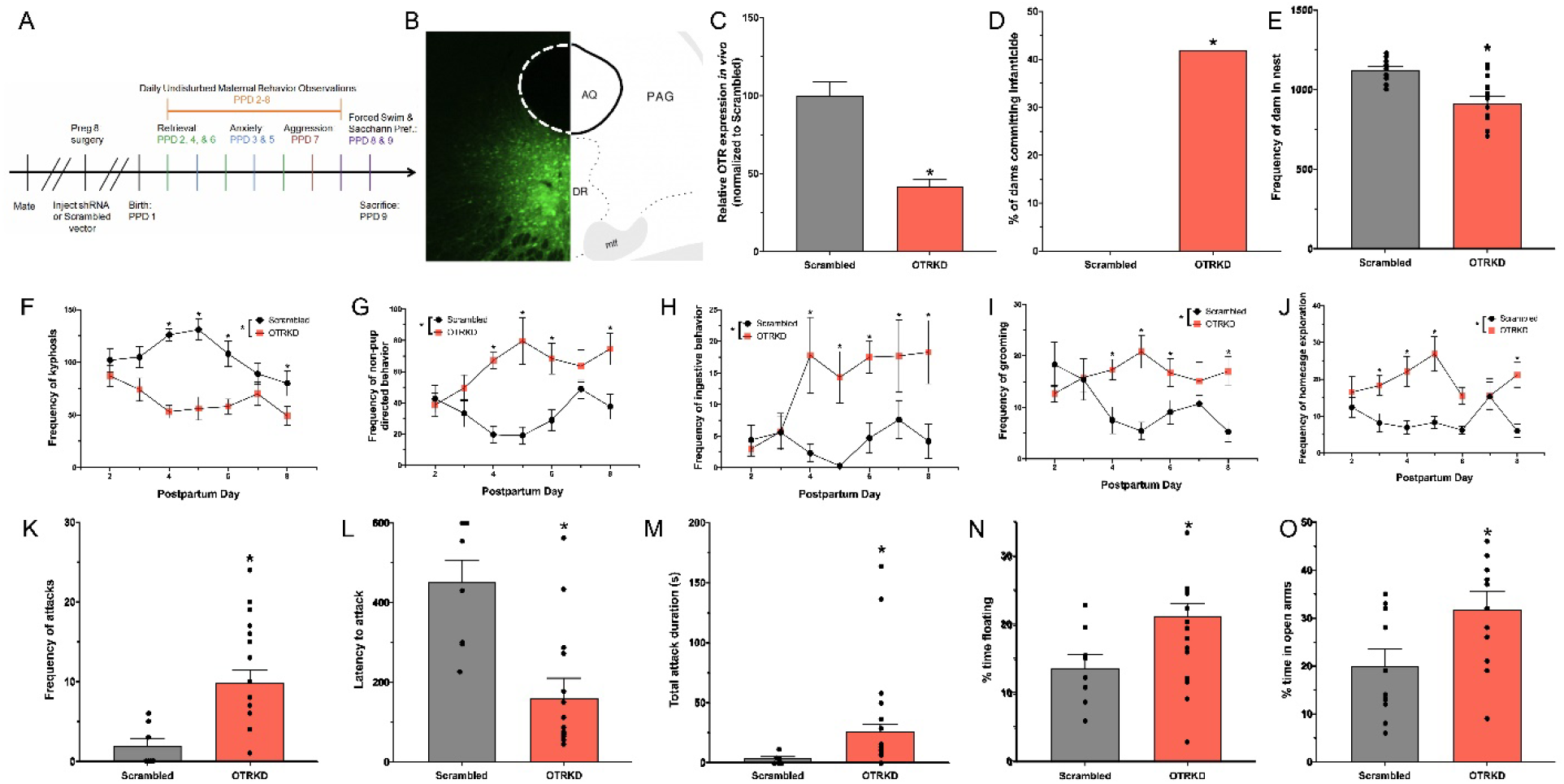
Midbrain OTRs regulate maternal socioemotional behaviors. *A*) Experimental timeline. *B*) Representative photomicrograph and brain atlas plates showing distribution of viral vector in the dorsal raphe (DR) and ventrolateral periaqueductal gray/DR lateral wing (PAGvl/DRlw). *C*) Oxytocin receptor (OTR) mRNA knockdown in scrambled-vector treated (Scrambled) and shRNA-vector treated (OTRKD) mother rats. *D*) Percentage of dams committing infanticide in scrambled and OTRKD mothers. Frequency of (*E*) mothers in the nest, (*F*) kyphotic nursing, (*G*) non-pup directed behavior, (*H*) ingestive behavior, (*I*) self-grooming, (*J*) home cage exploration, and (*K*) attacks by scrambled control and OTRKD dams. *L*) Latency to attack and (*M*) total duration of attacks by scrambled and OTRKD dams. Percentage of time (*N*) floating in the forced swim test and (*O*) in the open arms of the elevated plus maze. * indicates *p* < 0.05.

OTR-knockdown did produce a number of notable deficits in the females’ postpartum behavior. In particular, two to three days after giving birth, five of the twelve OTR-knockdown mothers were infanticidal compared to none of the controls (Fig 3D). Infanticide is extremely rare in primiparous female rats and somatosensory experience with pups can prevent infanticide in nulliparous rats^48^. This suggests that oxytocin signaling in the DR may be critical for how offspring cues prevent infanticide in the early postpartum period. Our results could have been due to decreased oxytocin activation of DR dopaminergic neurons, as activation of these neurons increases social investigation^43^. Consistent with this suggestion, OTR-knockdown also caused our mothers to spend less time with pups in the nest compared to the controls (Fig 3E).

In addition to promoting infanticide, OTR-knockdown decreased kyphotic (i.e., arched-back) nursing (Fig 3F), which is consistent with other work showing that global OTR blockade decreases kyphosis^49^. Instead of nursing, OTR-knockdown mothers displayed more non-pup directed behaviors including eating/drinking, self-grooming, and exploring the home cage (Figs 3G-J). The reduction in kyphosis after OTR knockdown may due to impaired oxytocin signaling in the PAGvl/DRlw given that lesioning this region of the brain greatly reduces kyphotic nursing in rats without affecting nursing in other postures^35^.

Our OTR knockdown also produced a >3-fold increase in the number of maternal attacks against male intruders and a >6-fold increase in the time that mothers spent attacking (Figs 3K-M). This pro-aggressive effect of OTR knockdown is consistent with studies showing that impairing DR serotonin cell function with selective lesions decreases maternal aggression^50^, that serotonin is positively associated with aggression in females^51^, and that destroying the PAGvl/DRlw doubles the frequency of maternal attacks on an intruder^35^.

Lastly, OTR knockdown increased the time that mothers spent floating in the forced-swim tests (an indicator of greater passive coping^52^) (Fig 3N) and decreased mothers’ anxiety-like behavior in an elevated plus maze (Fig 3O). Oxytocin typically has antidepressant effects^53^, so higher passive coping in the OTR knockdown mothers was not surprising. However oxytocin is a potent anxiolytic in rodents^54^, including postpartum female rats^55,56^, so the lower anxiety behavior in the knockdown mothers was unexpected. Oxytocin-facilitated serotonin release from the DR into the bed nucleus of the stria terminalis has recently been postulated to be part of an anxiety-<u>promoting</u> network^57^, though, which would be consistent with our results and suggest that oxytocin’s effects on anxiety are brain-region specific. Overall, our results demonstrate that high OTR expression in the DR and PAGvl/DRlw is critical for the normal display of many maternal socioemotional behaviors (Supplementary Table 2).

## OTRs influence cortical serotonin and perineuronal nets

It is interesting that a number of behavioral effects from our OTR knockdown did not emerge until postpartum days 2 or 3 (Figs 3D, F-J). One explanation for this is that OTRs in the DR and PAGvl/DRlw are involved in the <u>experience-dependent</u> plasticity underlying how touch regulates socioemotional behaviors. Reproduction is associated with tremendous neuroplasticity in the female mammalian cortex^58–64^. As examples, the primary auditory cortex (A1) undergoes changes in excitatory-inhibitory balance after female mice give birth^60,62–64^, and the volume of the primary somatosensory cortex (S1) dedicated to the ventrum almost doubles in nursing female rats^58,59^. Women also undergo cortical expansion or contraction as a result of motherhood^65,66^. These neuroplastic changes are necessary for mothers positive responses to offspring cues^58–64^, and are mediated by both maternal experience interacting with the young and hormones of reproduction^58,59,63,64^. Oxytocin is involved in this neuroplastic change. For example, oxytocin enhances cortical plasticity in the A1 by increasing responses to pup calls, which facilitates pup retrieval^60^, but it is unknown whether oxytocin affects the capacity for plasticity in other cortical regions. The primary somatosensory cortex (S1) and lateral orbitofrontal cortex (lOFC) transmit the affective content of somatosensory inputs^67–69^ and cortical plasticity requires serotonin^70^. Thus, OTRs in the DR may contribute to cortical plasticity in mothers.

We hypothesized that knocking down OTRs in the DR and adjacent PAGvl/DRlw disrupt the serotonin innervation of the sensory and orbitofrontal cortex, subsequently reducing the capacity for maternal experience-related neuroplasticity and thereby disrupting postpartum behaviors. To begin testing this, we used serotonin-immunoreactive fiber length as an indicator of the capacity for serotonin release^71,72^ and found that OTR knockdown did decrease serotonergic-immunoreactive fiber length in S1, but increased it in the lOFC (Figs 4D-F, 4O). OTR knockdown did not affect serotonin-immunoreactive fiber length everywhere in the cortex, though, as no differences between groups were found in the primary motor cortex (data not shown).

**Figure 4:**
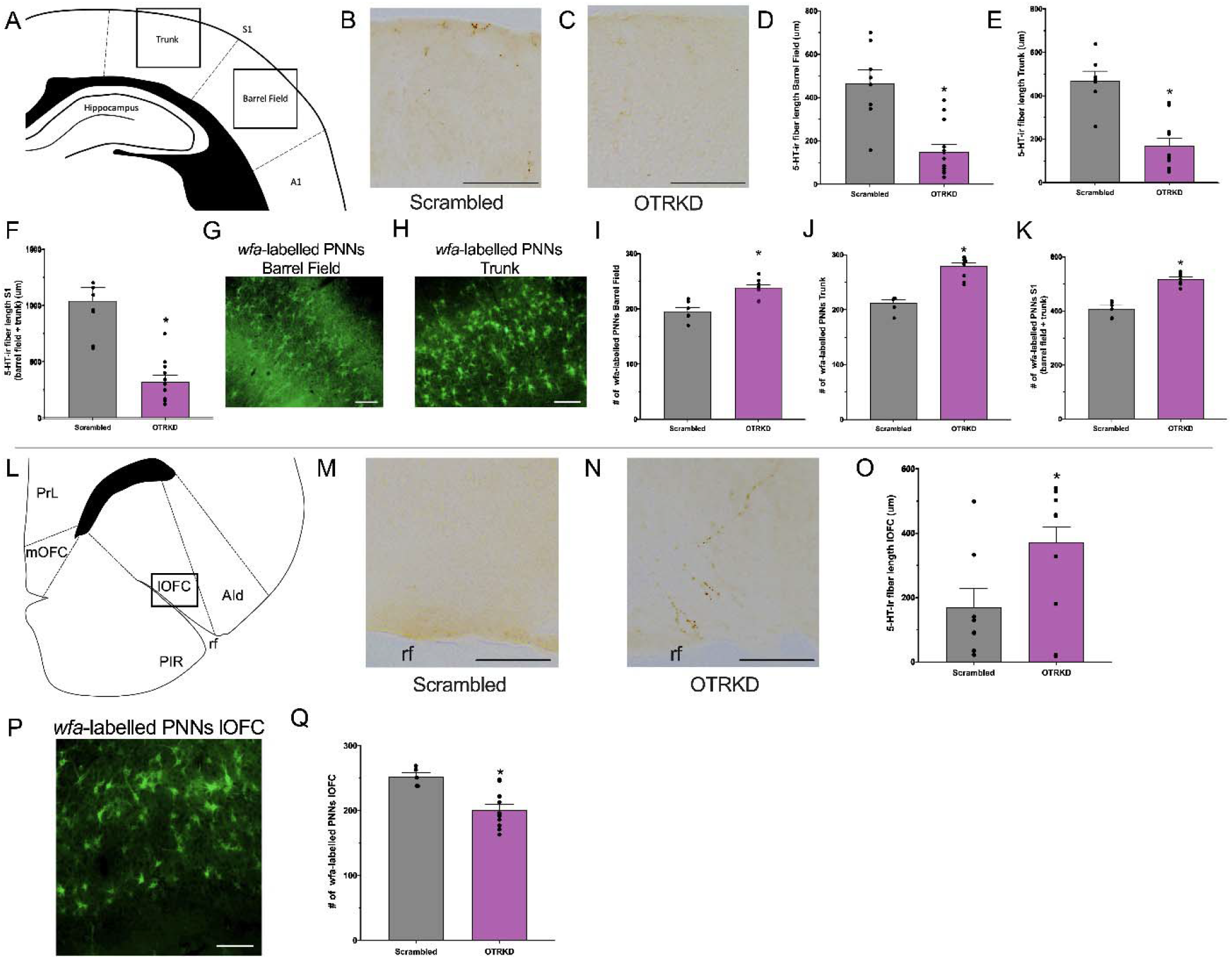
OTR knockdown affects cortical serotonin immunoreactivity and number of perineuronal nets. Atlas representations of areas of the (*A*) primary somatosensory cortex (S1) and (*L*) lateral orbitofrontal cortex (lOFC) analyzed^91^. Representative photomicrographs of serotonin-immunoreactive fibers in the (*B, C*) S1 and (*M, N*) lOFC of scrambled-vector treated (Scrambled) and shRNA-vector treated (OTRKD) mothers. Serotonin-immunoreactive fiber length in the (*D*) barrel field, (*E*) Trunk, (*F*) S1 (barrel field and trunk), and (*O*) lOFC in Scrambled control and OTRKD mother rats. Representative photomicrographs of perineuronal nets (PNNs) in the (*G, H*) S1 and (*P*) lOFC of Scrambled control and OTRKD mother rats. Number of PNNs in the (*I*) barrel field, (*J*) Trunk, (*K*) S1, and (*Q*) lOFC. * indicates *p* < 0.05. A1 = primary auditory cortex, AId = dorsal anterior insular cortex, mOFC = medial orbitofrontal cortex, PIR = piriform cortex, PrL = prelimbic cortex, rf = rhinal fissure, S1 = primary somatosensory cortex, SII = secondary somatosensory cortex. Scale bars in B, C, M and N = 100 μm.

We then used the number of perineuronal nets in the cortex as an indicator of mothers’ capacity for neuroplasticity. Perineuronal nets are extracellular matrix structures that surround the somata and proximal dendrites of GABAergic interneurons in the mature cortex^73^, and act as physical and chemical barriers to experience-dependent plasticity^73–75^. Given that greater serotonin release is negatively associated with the number of perineuronal nets in the medial prefrontal cortex and hippocampus^76,77^, and OTR knockdown decreased serotonin in the S1, we predicted that OTR-knockdown should increase PNNs in the S1. Likewise, since OTR KD increased serotonin in the lOFC, then OTR KD should decrease PNNs n the lOFC. OTR knockdown increased the number of perineuronal nets in the S1 (Figs 4I-K), while it decreased the number of perineuronal nets in the lOFC (Fig 4Q). These effects of OTR knockdown may have been driven by serotonin’s potentiation of cortical inhibition^68,78^, with serotonin release associated with decreased activity of the S1 in response to tactile stimulation in both female rats^79,80^ and women^68^. Perineuronal nets are generated in response to neural activity^73^, so greater S1 activity and lower lOFC activity could have led to the higher and lower number of perineuronal nets in the S1 and lOFC, respectively (Figs 4I-K, 4Q).

## Concluding remarks

Social touch is an evolutionarily conserved feature of positive behavioral interactions and bonding across animals of many species^1–5^. Using a mother-offspring model of social touch, we examined whether motherhood was associated with changes in aspects of the oxytocin system in the midbrain DR and PAGvl/DRlw, and whether OTRs there mediate maternal behavioral responses to social touch. We found that mothers’ responses to young and their affective behaviors were derailed by the loss of these midbrain OTRs (Fig 5). The work presented here further suggests that midbrain OTRs regulate socioemotional responses to social touch through long-term structural and activity-dependent changes in the S1 and lOFC. Future research should determine whether midbrain oxytocin signaling, particularly through DR serotonin cells, also mediate socioemotional consequences of gentle touch in other relationships such as between fathers and offspring, among romantically or sexually bonded adults, and in platonic relationships.

**Figure 5:**
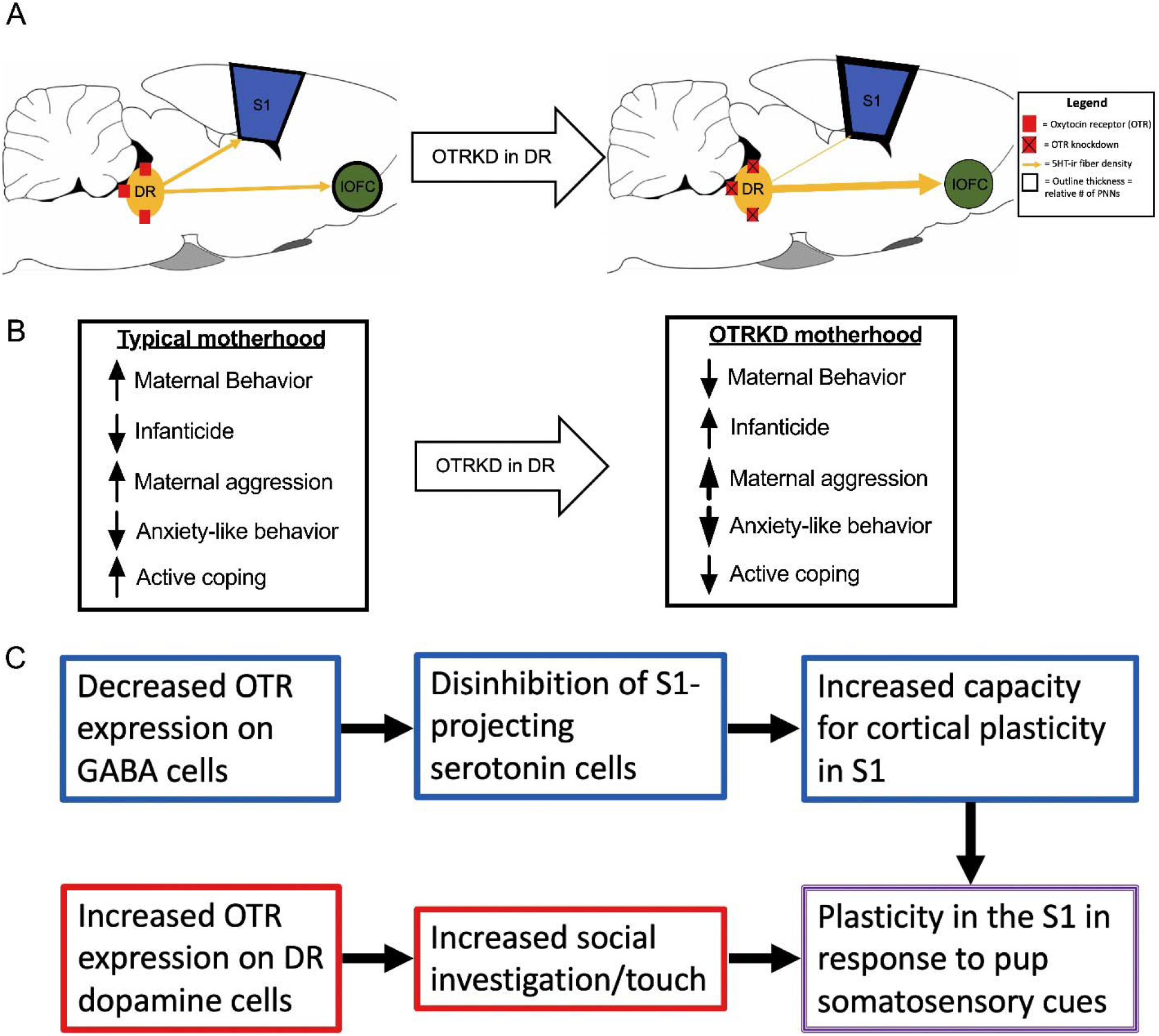
Summary of behavioral findings and proposed effects of oxytocin receptor knockdown (OTRKD). *A)* Oxytocin sensitive cells in the dorsal raphe (DR) project to the primary somatosensory cortex (S1) and the lateral orbitofrontal cortex (lOFC), which when disrupted by viral-mediated knockdown (OTRKD), leads to decreased serotonergic fiber density and increased perineuronal nets (PNNs) in the S1 and increased serotonergic fiber density and decreased PNNs in the lOFC. *B)* Motherhood leads to numerous behavioral changes, which are all disrupted after OTRKD in the DR. *C*) Proposed model for how the neurochemical changes in the DR of reproductive female rats leads to experience-dependent cortical plasticity.

## Methods

### Subjects

Female Long-Evans rats were descended from rats purchased from Harlan Laboratories (Indianapolis, IN), born and raised in the Lonstein colony at Michigan State University. Females were housed with 2 or 3 same-sex littermates in polypropylene cages (48 cm × 28 cm × 16 cm) containing wood chip bedding, food (Tekland rat chow) and water *ad libitum*, the room maintained on a 12:12 light/dark cycle (lights on 0700 hr), and temperature 22 ± 1 °C. After 65 days of age, estrus cycles were monitored by vaginal smearing. On a day of proestrus, subjects in reproductive groups were mated with a male from the colony and pregnancy confirmed by the presence of semen or vaginal plug. Subjects were singly housed 5-7 days before expected parturition and litters were culled to 8 pups (4M/4F). Females for behavioral analysis were group housed until stereotaxic surgeries on pregnancy day 8, then singly housed. All procedures were performed in accordance with the NIH Guide for Care and Use of Laboratory Animals and approved by the Michigan State University IACUC.

### Oxytocin receptor autoradiography

On a scheduled day of sacrifice (diestrous (DV), pregnancy day 10, parturition - within 3 hours after delivery of last pup, or postpartum day 7; *n*=5/group) females were rendered unconscious with CO_2_, and decapitated. Brains were frozen with isopentane and stored at −80°C until sectioning. Brains were cut coronally into 15-μm sections and mounted in 4 series using a cryostat (Leica CM1950, Nussloch, Germany) to obtain 16 matching sections containing the DR (−7.3 to −8.7 mm from bregma) and 14 matching sections including PAGvl (−7.1 to −8.3 mm from bregma). Slides were stored at −80 °C and on a day of autoradiography allowed to thaw at room temperature. Slides were then incubated for 2 mins in 1% paraformaldehyde, followed by two rinses for 10 mins each in 50 mM Tris-HCl. Slides were incubated for 1 hr in 50 mM Tris-HCl containing 0.1% bovine serum albumin and 50 pM of the radioactive tracer ornithine vasotocin analog, [125I]-OVTA (Perkin-Elmer; # NEX254050UC). Slides were rinsed twice in ice-cold 50 mM Tris-HCl for 5 mins, and a final wash in ice-cold water. Slides dried overnight at 4 °C and placed against Kodak BioMaxMR film for 7 days and developed using a Kodak X-OMAT 1000A Processor (Kodak, Rochester, NY). Optical density for each section was analyzed using ImageJ (NIH, Bethesda, MD). Mean density per region was calculated by subtracting background signal from the density of the region of interest. Data from two adjacent brain sections were combined into a single data point for each animal. The rostral DR data were from two sections between −7.3 to −7.6 mm from bregma; the medial DR from four sections between −7.7 to −8.3 mm; and the caudal DR from two sections between −8.5 to −8.7 mm as defined by Lowry and colleagues^18^.

### Oxytocin fiber immunohistochemistry

Subjects were overdosed with sodium pentobarbital either on diestrus, within three hours of birth, or postpartum day 7 (*n*=5/group). Subjects were perfused with saline followed by 4% paraformaldehyde, brains postfixed overnight, and submerged in 30%. Tissue was cut into 40-μm sections in four series and stored in cryoprotectant at −20°C. Seven matched sections containing the DR (−7.1 to −8.9 mm from bregma) and four matched sections containing the PAGvl (−7.3 to −8.3 mm from bregma) were analyzed. All rinses were in TBS. Sections were incubated in 0.1% sodium borohydride for 10 min, followed by a 10-min incubation in a 1% H_2_O2 in 0.3% triton-TBS. Sections were blocked in 20% NGS and 0.3% Triton-X for 60 min. Sections were incubated in a triton-TBS containing 2% NGS and a mouse anti-oxytocin polyclonal antiserum (MAB5296, Millipore, Burlington, MA; 1:300) for 72 hrs at 4 °C, then in biotinylated goat anti-mouse secondary antiserum (BA-2000; Vector Labs, Burlingame, CA; 1:500) in a triton-TBS solution containing 2% NGS for 1 hr at room temperature. The sections were then incubated in ABC (PK 6100, Vectors Laboratories, Burlingame, CA) for 1 hr and oxytocin-ir visualized using Vector-SG (SK-4700; Vector Laboratories). Sections were mounted onto microscope slides and coverslipped. The length of oxytocin-ir fibers in the entire area of the brain structures was traced on each section by experimenter’s blind to the subjects’ experimental condition under 200X magnification. The length of oxytocin-ir fibers traced across all sections per site was used for data analyses. Additionally, the DR was divided into its rostral (two sections between −7.1 to −7.3 mm from bregma); medial (three sections between −7.6 to −8.3 mm); and caudal (two sections between −8.5 to −8.9 mm) subregions for follow-up analyses.

### Oxytocin receptor and tyrosine hydroxylase or tryptophan hydroxylase double-label immunohistochemistry

Subjects were overdosed with sodium pentobarbital either on diestrus or within 3 hours after birth (*n*=5/group). Subjects were perfused with saline followed by 4% paraformaldehyde, brains postfixed overnight, and submerged in 30%. Tissue was cut into 40-μm sections in four series and stored in cryoprotectant at −20°C. Dual label immunohistochemistry was used to examine the percentage of DR tyrosine hydroxylase (TH) and tryptophan hydroxylase (TPH) immunoreactive cells also expressing OTR immunoreactivity. All rinses were in PBS. Slides were removed from −80 °C to reach room temperature and slides for TH-OTR immunohistochemistry blocked in 0.2% triton-PBS solution containing 5% NDS for 3 hrs. Slides were incubated in the highly purified version of the rabbit anti-OTR primary antiserum generated by and gifted to us fromDrs. Robert Froemke and Moses Chao at NYU (1:200) and mouse anti-TH primary antiserum (MAB5986, Millipore, Burlington, MA; 1:2000) for 48 hrs at 4 °C. Slides were incubated in donkey anti-mouse Alexafluor 647 (A21448, Fisher Scientific, Pittsburgh, PA; 1:500) and donkey anti-rabbit Alexafluor 488 (A21206, Fisher Scientific; 1:500) secondary antisera for 2 hrs, followed by coverslipping using Fluoromount G (Southern Biotech, Birmingham, AL).

Slides for TPH-OTR immunohistochemistry were blocked in 5% NDS in PBS for 1 hr and then incubated in sheep anti-TPH primary antiserum (T857501VL, Sigma-Aldrich, St. Louis, MO; 1:1200) overnight at room temperature. Slides were then blocked for 3 hrs in a 0.2% triton-PBS solution containing 5% NDS, then in the purified rabbit anti-OTR primary antiserum for 48 hrs at 4 °C. Slides were then incubated in donkey anti-sheep Alexafluor 647 (A21448, Fisher Scientific; 1:500) and donkey anti-rabbit Alexafluor 488 (A21206, Fisher Scientific; 1:500) secondary antisera for 90 mins, followed by coverslipping using Fluoromount G.

Sections were analyzed with a Nikon C2 confocal microscope, using 488 nm (OTR immunofluorescence) and 647 nm (TH/TPH immunofluorescence) lasers and band pass emission filters for wavelength selection. High-resolution confocal fluorescence was collected through a single, variable pinhole aperture and recorded using three high-sensitivity photomultiplier (PMT) detectors. Z-stack images were collected at 1.64-μm intervals through three sections containing the rostral DR (location of TH-ir neurons) and seven sections containing the entire DR (location of TPH-ir neurons) per animal. Colocalization with of OTR-ir with TH/TPH-ir was determined via 3D reconstruction using a Z-stack orthogonal viewer.

### Tyrosine hydroxylase immunohistochemistry

Subjects were overdosed with sodium pentobarbital either on diestrus, within three hours of giving birth, postpartum day 7, or postpartum day 18 (*n* = 6/group). Subjects were perfused with saline followed by 4% paraformaldehyde, brains postfixed overnight, and submerged in 30%. Tissue was cut into 40-μm sections in four series and stored in cryoprotectant at −20°C. We performed quantitative histological analysis as previously described^81^. Five matched sections per subject containing the DR (−7.0 to −7.8 mm from bregma) were analyzed. To confirm that differences among groups in TH-ir neuron number were specific to the DR, the number of TH-ir cells was also analyzed in three matched sections per subject containing the VTA (−5.65 to −6.06 mm from bregma). The number of TH-ir cells in the DR (complete visual area) was counted on each section by naive experimenters under 200X magnification. Somata with any visible TH-ir were quantified. The summed number of TH-ir cells in all sections was used for data analyses. In addition, the percentage of total DR area covered by TH-ir pixels (using standardized light level and threshold for optic density with NIS Elements) was also analyzed.

### In situ hybridization analysis of oxytocin receptor and tyrosine hydroxylase, tryptophan hydroxylase, or glutamic acid decarboxylase mRNA in the DR

On a scheduled day of sacrifice (diestrous or within three hours of birth; *n*=3/group) females were rendered unconscious with CO_2_, and decapitated. Brains were frozen with isopentane and stored at −80°C until sectioning. Brains were cut coronally into 20-μm sections into 16 series through the DR. This resulted in 7 sections through the DR (2 sections containing the rostral DR subregion, 3 sections containing the medial subregion, and 2 sections containing the caudal subregion). Slides were stored in −80 °C until RNAscope® processing. On the day of RNAscope® processing to detect OTR and TH, TPH, or glutamic acid decarboxylase (GAD) colocalized hybridization, slides were removed from −80 °C and immersed in 4°C 4% paraformaldehyde for 15 min. Tissue was dehydrated through serial ethanol solutions dried for 5 min. Slides were then incubated with ~5 drops of RNAscope protease IV (Cat. 322340) for 30 min and rinsed twice with 1 X PBS. Probes were warmed for 10 min at 40 °C, cooled to room temperature, then mixed for their respective runs: Run 1) GAD (Cat. 316401), OTR (Cat. 483671), and TPH (Cat. 316411); Run 2) TH (314651) and OTR. Excess liquid was removed from the slides and ~4-5 drops of the mixed probes were added. Slides on the HybEZ™ humidity control tray were then placed into the HybEZ™ oven for 2 hrs at 40 °C. After every incubation, slides were rinsed 2 times for 2 minutes in wash buffer. Slides were incubated in ~4-5 drops of Amp 1-FL for 30 min at 40 °C in the HybEZ™ oven, then in Amp 2-FL for 15 min at 40 °C. Following rinses, the slides were then incubated in Amp 3-FL for 30 min at 40 °C, followed by Amp 4-FL for 15 min at 40 °C. Slides were then incubated with DAPI (~ 30 sec), followed by ~2 – 3 drops of Prolong Gold Antifade Mountant (P36930, Thermo Fisher Scientific, Waltham, MA) and coverslipped. Sections were analyzed with a Nikon C2 confocal microscope, using 488 nm (GAD/TH fluorescence), 568 nm (OTR fluorescence), and 647 nm (TPH fluorescence) lasers and band pass emission filters for wavelength selection. Two sections containing the rostral dorsal raphe (for colocalization of TH/OTR) and seven sections containing the entire dorsal raphe (for colocalization of TPH/OTR and GAD/OTR) per animal were analyzed as described above.

### Creation of viral construct

Candidate shRNAs against OTR mRNA (NCBI reference sequence: NM_012871.3) were designed using previously validated methods^82^ and cloned into an adeno-associated virus (AAV) genome under control of the H1 promotor. The same genome also carried a GFP reporter under the control of the chicken beta-actin promotor/cytomegalovirus enhancer promotor hybrid (pCBA)^82^. shRNA candidates were evaluated *in vitro* using a dual-luciferase (Firefly-Renilla) assay system, and the best candidate (shRNA sequence CGGTGAAGATGACCTTCAT) produced ~79% knockdown of OTR mRNA (Supplementary Fig 1A). Scrambled shRNA was used as a negative control. Vector genomes were packaged into an AAV 9 capsid, which produces high transduction of the DR^83^, using a standard triple transfection protocol followed by recovery of vector from media and cells^84,85^.

### Stereotaxic injections

On pregnancy day 8, rats were anesthetized with ketamine (90 mg/kg IP; Butler, Dublin, OH) and xylazine (8 mg/kg IP; Butler, Dublin, OH) and placed in a stereotaxic apparatus. A hole was drilled into the skull above the DR (A/P=7.8 mm, M/L=0.0 mm from bregma) and 1 μL of OTR-shRNA or the scrambled control vector slowly injected (~0.5 μL/ 5 mins) through a Neuros Hamilton syringe at 6.7 mm ventral from the skull (*n* =10-12/group for behavioral analysis). The needle remained for 10 min and then slowly retracted. Subjects received twice-daily subcutaneous injections of buprenorphine (0.015 mg/kg) for 1 day after surgery and then left undisturbed until parturition.

### In vivo determination of oxytocin receptor knockdown

Before any behavioral studies were conducted, level of oxytocin receptor knockdown was determined *in vivo*. Within 3 hrs of birth, OTRKD- and Scramble-injected dams used to determine level of oxytocin receptor knockdown (*n* = 6/group) were rendered unconscious with CO_2_ and decapitated. Brains were removed, flash frozen with isopentane, and stored at −80 °C. Brains were cut coronally into 300-μm-thick sections to obtain 3 sections that included the DR (−7.3 to −8.3 mm from bregma). The DR was punched from the sections using a 1-mm-diameter micropuncher (Harris Micropunch, Hatfield, PA) to examine OTR mRNA levels as previously done^86^. Because the vasopressin V1a receptor gene has high homology with the oxytocin receptor gene, PCR was also run for V1a receptor to confirm specificity of our viral construct. PCR products of each primer set were sequenced at the RTSF Genomics Core at Michigan State University to confirm specificity. The ΔΔCT method was used to calculate fold change between groups, with OTR and V1a normalized to HPRT-1^87^.

### Undisturbed maternal behavior observations

Dams’ behavior in the homecage was recorded 3 times daily (0900, 1300, and 1500 hr) for 30 min each on postpartum days 2 – 8. These timepoints ensured that we would observe maternal caregiving, given that dams spend most of their time interacting with their litters during the light photophase^88^. Spot checks were made every 30 s and maternal behaviors recorded included pup licking, nest building, hovering over the pups, and nursing in three distinct postures (kyphosis/crouched, supine/on side, and prone/flat). The four in-nest postures were analyzed individually as well as in two broader categories – those involving erect postures (hovering over and kyphosis/crouched) and those involving passive posture (supine and prone). Non-maternal behaviors included sleeping away from the pups, exploring, self-grooming, and eating or drinking. Quality of the dams’ nests were scored during each undisturbed observation on a scale of 0-3 (0=no nest, 1=poor nest, 2=partial nest, 3=complete nest with high walls). Frequencies of each behavior were totaled for each day and those totals used for data analyses. The multiple observers established an inter-rater reliability of >90% before beginning data collection.

### Retrieval testing

Litters were briefly removed from the nest to induce maternal as an indicator of maternal motivation under mildly stressful conditions. On postpartum days 2, 4, and 6, immediately follow undisturbed maternal behavior observations, litters removed and placed in an incubator set to nest temperature (34 °C) for 15 min. The pups were then scattered in the home cage opposite the nest site. Latencies for the dams to retrieve each pup and hover over all of the pups in the nest were recorded. Maternal behaviors were then recorded by spot checks every 30 s for an additional 10 min. Given that there were no main effects of postpartum day, nor any interactions between postpartum day and group on any behaviors, the behavioral results from the three retrieval days were averaged together for analyses.

### Elevated plus maze and light-dark box texting

On postpartum day 3, following undisturbed maternal behavior observations (~1600-1700 hr), dams were brought in their homecage to a nearby room containing an elevated plus-maze and tested as previously described^89^. The recording of the behavior was scored with a computerized data acquisition system that allowed recording the time spent in the open arms and closed arms, and the frequency of each. An entry into an arm was coded when the dam placed her head and both paws into an arm, while time spent in the center square was recorded as time spent in neither arm.

On postpartum day 5, following undisturbed maternal behavior observations (~1600-1700 hr), dams were brought in their homecage to a nearby behavior testing room containing the light-dark box and tested as previously described^90^. Females’ latency to enter the dark chamber was scored, total time spent, entries, and nose pokes into the light chamber was then scored using the recording and a computerized data acquisition system. Rears while in the light chamber were also scored.

### Maternal aggression

On postpartum day 7, following undisturbed maternal behavior observations, dams were brought in their homecage to a nearby behavior testing room and a male intruder was placed in their cage as previously described^50^. Following the 10-min test, males were removed from the cage and sacrificed by CO_2_ asphyxiation. Females were then returned to the colony room.

### Saccharin preference test

A saccharin habituation period began in the morning of postpartum day 8 (~0800 hr), when two water bottles (one containing 0.1% saccharin and another water) were weighed and placed in the home cage. Eight hr later, the bottles were weighed again and their positions switched to avoid side preference confounds. Testing on the morning of postpartum day 9 (~0800 hr) involved both bottles being weighed, and then removed from the cage along with food for 4 hr. Following deprivation, food and both water bottles were returned to the home cage and pups removed to limit potential distraction. Following a 1 hr testing period, the bottles were removed and weighed, and the regular water bottle returned to the cage. The change in bottle weights between pre- and post-1 hr testing was used for data analyses.

### Forced swim test

Following undisturbed behavior observation on postpartum day 8, dams were brought to a nearby testing room containing a 50 cm × 20 cm Plexiglas cylinder 40 cm full of 24°C water for pre-exposure. Dams were placed in the cylinders and behavior recorded for 15 min. Dams were then removed from the cylinder, towel-dried, and placed into a new cage containing a prewarmed heating pad. Once dry, dams were returned to their homecages. The next day, 1 hr after saccharin preference testing the dams were brought back to the behavior testing room and placed into the cylinder for 10 min. After removal and drying, dams were placed in the homecage until sacrifice that evening. Females’ behavior was then scored from the digital recording with a computerized data acquisition system (Soloman Coder). The percentages of time swimming and floating were coded and analyzed.

### Sacrifice, perfusion, and brain extraction

Following forced-swim testing on postpartum day 9, subjects were perfused with saline followed by 4% paraformaldehyde, the brains extracted, postfixed overnight, and submerged in 30% sucrose. Brains were cut into 40-μm sections in three series and stored at −20 °C in a sucrose-based cryoprotectant. One series was processed to determine viral injection sites and another series for serotonin-immunoreactive fiber analysis in the forebrain.

### GFP immunohistochemistry for injection localization

All rinses were in PBST. Sections were rinsed in PBST and then blocked in a 0.1% triton-PBS solution containing 2% NDS for 1 hr at room temperature. Sections were then incubated in purified rabbit anti-GFP primary antiserum (A6455; Thermo-Scientific, Waltham, MA; 1:10,000) for 24 hrs at 4 °C, then in donkey anti-rabbit Alexafluor 488 secondary antisera (A21206, Fisher Scientific, Pittsburgh, PA; 1:500) for 2 hrs at room temperature, followed by mounting and coverslipping using Fluoromount-G. The entirety of the dorsomedial tegmentum was scanned for GFP-ir cells under 40× magnification. Any subjects that had GFP-ir cells outside the dorsomedial tegmentum were removed from further analyses.

### Serotonin immunohistochemistry

To help verify that shRNA infusion did not produce gross impairments in DR cell viability, three matched sections per subject containing the DR (−7.5 to −8.4 mm from bregma) were selected for analysis of the number of serotonin-ir cells. Additionally, to determine the effects of OTR knockdown on serotonin-ir fiber length in cortical sites relevant to maternal behavior, one section containing each of the S1 (−2.85 mm from bregma), the motor cortex (M1; −0.0 mm from bregma), and the lOFC (+3.2 mm from bregma) were analyzed (*n* = 6/group).

Immunohistochemistry was conducted using methods previously described^50^. The number of serotonin-ir cells in the DR (complete visual area) was counted on each section by naive experimenters under 100X magnification. Somata with any visible serotonin immunoreactivity were quantified. The summed number of serotonin-ir cells in all sections for each subject was used for data analyses. The length of serotonin-ir fibers in the trunk and barrel cortex regions of the somatosensory cortex, motor cortex, and lOFC were traced bilaterally under 200X magnification. The length of serotonin-ir fibers traced per site for each subject was used for data analyses.

### Perineuronal net assay

Perfusion-fixed tissue was rinsed 5 times in PBS for 5 min each, then incubated in fluorescently tagged-*wisteria floribunda lectin* (Vector Laboratories; Cat. No. FL-1351; 1:200) for 2 hrs at room temperature in the dark. Tissue was then rinsed 5 times in PBS for 5 min, then mounted and coverslipped using Fluormount-G. Images of the S1, lOFC, and M1 were taken at 10 x magnification under epilluminescence.

### Statistical analyses

Optical density of OTR binding (Experiment 1) and oxytocin-ir fiber length (Experiment 2) were analyzed with repeated-measures ANOVAs (with Greenhouse-Geisser correction) involving rostrocaudal levels as the repeated measure and reproductive state as the between-subject variable. The number of TH-ir neurons in the DR across reproduction (Experiment 3b), were analyzed with one-way ANOVAs. In cases of statistical significance, LSD post-hoc tests were conducted to compare across rostrocaudal level and between groups. For the immunohistochemical (Experiment 3a) and *in situ* hybridization (Experiment 4) colocalization studies involving two groups, Student’s *t*-tests were used. All data were normally distributed with no significant outliers using Dixon-Q extreme outlier tests. *p* < 0.05 was considered statistically significant.

Undisturbed maternal behavior was analyzed using mixed-design repeated-measures ANOVAs (with Greenhouse-Geisser correction) involving time (i.e., postpartum day) as the repeated measure and experimental condition (i.e., scrambled or oxytocin receptor shRNA) as the between-subject variable. Data for all between-subjects variables were normally distributed. Analysis of infanticide occurrence was done using Fisher’s exact test. Two dams completely cannibalized their litters and the replacement litters given to them, so were removed from all behavioral analyses other than infanticide. Another three dams killed 1 - 3 pups in their litters, but after providing replacement pups, these dams were able to maintain the litter and remained in the study. Retrieval, aggressive, anxiety-like, and depressive-like behaviors were analyzed using Student’s *t*-tests. In cases of unequal variances, Welch’s *t*-tests were used. *p* < 0.05 was considered statistically significant.

## End notes

## Acknowledgements

The authors would like to thank E. Vitale, M. Davis, M. Ahern and Katrina Linning for the help collecting behavioral data. Dr. K. Krishnan for her input regarding perineuronal nets and cortical plasticity. Drs. R. Froemke and M. Chao for the gifted OTR primary antibody. Dr. A. Veenema for her assistance with *in situ* hybridization. These studies were funded by NICHD HD097085 awarded to JSL.

## Author contributions

ZAG conducted all experiments and data analyses, along with writing and editing the manuscript. EGF conducted behavioral analysis and serotonin fiber and perineuronal net analyses, along with editing the manuscript. FPM designed and produced of the OTR shRNA vector, along with editing the manuscript. JSL helped with experimental design and data analyses, along with writing and editing the manuscript.

