## Supplementary Figure 1 for "Dorsal raphe oxytocin receptors regulate the neurobehavioral consequences of social touch"

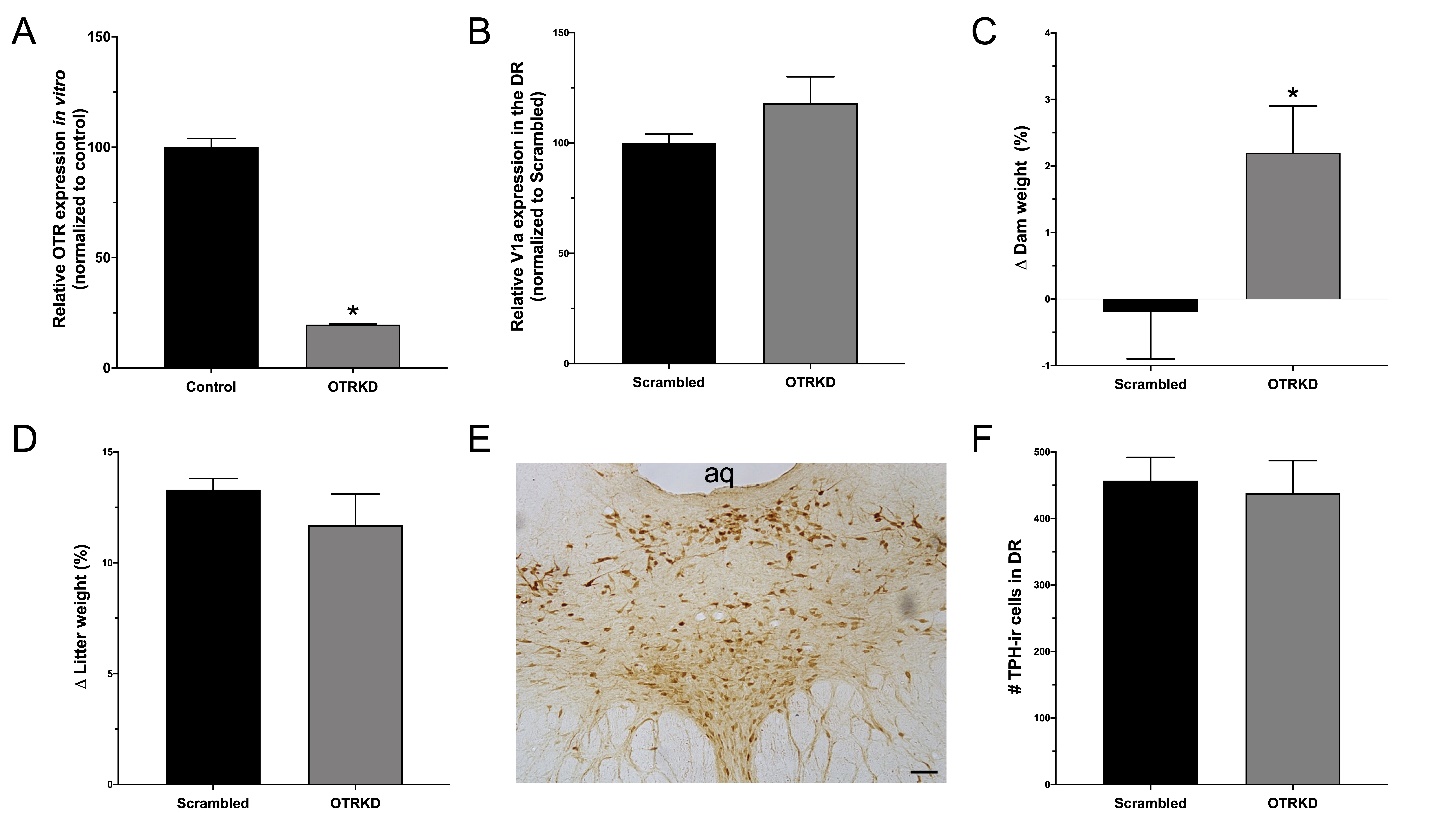

*Supplementary Figure 1:* **OTRKD did not generally affect maternal or litter health, or dorsal raphe serotonin cell number.** *A*) *In vitro* oxytocin receptor expression in control and shRNA-vector treated (OTRKD) cells. *B*) Vasopressin V1a receptor expression *in vivo* in scrambled-treated (Scrambled) and OTRKD dams. Change in (*C*) dam weight and (*D*) litter weight after Scrambled and OTRKD treatment. *E*) Representative photomicrograph of tryptophan hydroxylase (TPH) immunoreactive cells in the dorsal raphe (DR). *F*) Number of TPH-ir cells in the DR of Scrambled and OTRKD dams.
