## Supplementary Table 3 for "Dorsal raphe oxytocin receptors regulate the neurobehavioral consequences of social touch"

| **Maternal Aggression** | **Scrambled (Mean ± SEM)** | **OTRKD (Mean ± SEM)** | ***t*_(20)_; *p*; *d*** |
| --- | --- | --- | --- |
| Attack latency (s) | 451 ± 56 | 159 ± 51 | 3.89; <0.01*; 1.84 |
| Attack frequency | 2 ± 1 | 10 ± 2 | 3.96; <0.01*; 1.95 |
| Average attack duration (s) | 1 ± 0.3 | 3 ± 1 | 2.65; 0.02*; 1.31 |
| Total attack duration (s) | 4 ± 1 | 26 ± 6 | 3.52; <0.01*; 1.58 |
| Frequency frontal attacks | 2 ± 1 | 7 ± 2 | 2.28; 0.04*; 1.12 |
| Frontal attack duration | 3 ± 1 | 20 ± 6 | 2.28; 0.04*; 1.15 |
| Frequency of lateral attacks | 1 ± 1 | 3 ± 1 | 2.52; 0.02*; 1.21 |
| Lateral attack duration | 1 ± 1 | 6 ± 2 | 2.52; 0.02*; 1.27 |
| **Depressive-like Behaviors** |  |  |  |
| Saccharin preference over water | 64 ± 9 | 78 ± 10 | 1.02; 0.32; 0.48 |
| % Time floating | 13.8 ± 2.0 | 21.3 ± 1.9 | 2.70; 0.02*; 1.28 |
| **Elevated Plus Maze** |  |  |  |
| % Entries into open arms | 40 ± 2 | 50 ± 2 | 3.11; <0.01*; 1.34 |
| % Time in open arms | 20 ± 4 | 32 ± 4 | 2.20; 0.04*; 0.95 |
| Total arm entries | 25 ± 3 | 37 ± 3 | 2.63; 0.02*;1.15 |
| Closed-arm entries | 15 ± 2 | 19 ± 2 | 1.25; 0.23; 0.55 |
| **Light-Dark Box** |  |  |  |
| Latency (s) to enter dark chamber | 6 ± 1 | 7 ± 2 | 0.49; 0.64; 0.23 |
| Duration (s) in light chamber | 28 ± 8 | 55 ± 11 | 1.90; 0.07; 0.84 |
| Frequency of transitions between chambers | 2 ± 1 | 4 ± 1 | 2.05; >0.05; 0.91 |
| Frequency of stretches in the light chamber | 10 ± 2 | 15 ± 2 | 2.13; <0.05*; 0.92 |
| Frequency of rears in the light chamber | 3 ± 1 | 4 ± 1 | 0.18; 0.86; 0.07 |

*Supplementary Table 3:* **Oxytocin receptor knockdown affected numerous maternal socioemotional behaviors.** Mothers’ behaviors (Mean ± SEM) displayed in midbrain scrambled and oxytocin receptor knockdown rats tested in an elevated plus maze, light-dark box, resident-intruder paradigm, saccharin preference test, the forced swim test. * indicates significant group difference.
