## Supplementary Table 1 for "Dorsal raphe oxytocin receptors regulate the neurobehavioral consequences of social touch"

| **Immunohistochemistry Colocalization** | Diestrous Virgins (M ± SEM) | Parturition (M ± SEM) | Group (*t_(6)_; p_)_* |
| --- | --- | --- | --- |
| **Total DR** |  |  |  |
| # TPH-ir cells | 680 ± 43 | 702 ± 79 | 0.25; 0.81 |
| # TPH-ir cells expressing OTR-ir | 225 ± 26 | 277 ± 39 | 1.10; 0.32 |
| %TPH-ir cells expressing OTR-ir | 33 ± 4 | 39 ± 3 | 1.10; 0.31 |
| **Rostral DR** |  |  |  |
| # TH-ir cells | 124 ± 7 | 238 ± 28 | 3.89; 0.051 |
| # TH-ir cells expressing OTR-ir | 53 ± 3 | 126 ± 12 | 5.76; 0.02* |
| %TH-ir cells expressing OTR-ir | 31 ± 1 | 39 ± 1 | 6.29; <0.01* |
| # TPH-ir cells | 188 ± 45 | 128 ± 26 | 1.25; 0.27 |
| # TPH-ir cells expressing OTR-ir | 52 ± 13 | 50 ± 9 | 0.13; 0.91 |
| %TPH-ir cells expressing OTR-ir | 28 ± 2 | 41 ± 5 | 2.00; 0.10 |
| **Medial DR** |  |  |  |
| # TPH-ir cells | 481 ± 29 | 483 ± 40 | 0.03; 0.98 |
| # TPH-ir cells expressing OTR-ir | 167 ± 34 | 177 ± 31 | 0.23; 0.83 |
| %TPH-ir cells expressing OTR-ir | 34 ± 5 | 36 ± 3 | 0.39; 0.71 |
| **Dorsomedial DR** |  |  |  |
| # TPH-ir cells | 198 ± 14 | 160 ± 16 | 1.82; 0.12 |
| # TPH-ir cells expressing OTR-ir | 68 ± 15 | 63 ± 13 | 0.28; 0.79 |
| %TPH-ir cells expressing OTR-ir | 34 ± 5 | 38 ± 4 | 0.68; 0.52 |
| **Ventromedial DR** |  |  |  |
| # TPH-ir cells | 188 ± 18 | 219 ± 17 | 1.24; 0.26 |
| # TPH-ir cells expressing OTR-ir | 65 ± 16 | 76 ± 13 | 0.52; 0.62 |
| %TPH-ir cells expressing OTR-ir | 33 ± 5 | 34 ± 4 | 0.14; 0.89 |
| **Caudal DR** |  |  |  |
| # TPH-ir cells | 77 ±12 | 122 ± 28 | 2.05; 0.11 |
| # TPH-ir cells expressing OTR-ir | 26 ± 3 | 66 ± 2 | 11.14; <0.001* |
| %TPH-ir cells expressing OTR-ir | 34 ± 3 | 57 ± 10 | 2.17; 0.10 |
| **PAGvl/DRlw** |  |  |  |
| # TPH-ir cells | 96 ± 2 | 104 ± 10 | 0.74; 0.51 |
| # TPH-ir cells expressing OTR-ir | 34 ± 6 | 39 ± 5 | 0.67; 0.53 |
| %TPH-ir cells expressing OTR-ir | 35 ± 6 | 37 ± 2 | 0.31; 0.7 |
| ***In situ* hybridization colocalization** | Diestrous Virgins (M ± SEM) | Parturition (M ± SEM) | Group (*t_(4)_; p_)_* |
| **Total DR** |  |  |  |
| # TPH cells | 951 ± 135 | 842 ± 100 | 0.65; 0.55 |
| # TPH cells expressing OTR | 262 ± 51 | 179 ± 44 | 1.23; 0.29 |
| %TPH cells expressing OTR | 27 ± 2 | 21 ± 3 | 1.94; 0.13 |
| # GAD^67^ cells | 830 ± 64 | 620 ± 43 | 2.71; 0.053 |
| # GAD^67^ cells expressing OTR | 318 ± 38 | 211 ± 32 | 2.15; 0.10 |
| % GAD^67^ cell expressing OTR | 38 ± 2 | 34 ± 3 | 1.19; 0.30 |
| **Rostral DR** |  |  |  |
| # TH cells | 86 ± 3 | 112 ± 15 | 1.67; 0.33 |
| # TH cells expressing OTR | 22 ± 5 | 40 ± 10 | 1.91; 0.15 |
| %TH cells expressing OTR | 25 ± 4 | 35 ± 4 | 1.47; 0.24 |
| # TPH cells | 220 ± 61 | 71 ± 22 | 2.30; 0.08 |
| # TPH cells expressing OTR | 74 ± 36 | 19 ± 7 | 1.53; 0.26 |
| %TPH cells expressing OTR | 30 ± 6 | 26 ± 4 | 0.68; 0.54 |
| # GAD^67^ cells | 199 ± 23 | 129 ± 4 | 2.97; 0.09 |
| # GAD^67^ cells expressing OTR | 92 ± 7 | 34 ± 7 | 6.18; <0.01* |
| % GAD^67^ cells expressing OTR | 47 ± 2 | 26 ± 4 | 4.29; 0.01* |
| **Medial DR** |  |  |  |
| # TPH cells | 639 ± 74 | 650 ± 58 | 0.12; 0.91 |
| # TPH cells expressing OTR | 164 ± 25 | 134 ± 33 | 0.72; 0.51 |
| %TPH cells expressing OTR | 26 ± 2 | 20 ± 3 | 1.31; 0.26 |
| # GAD^67^ cells | 430 ± 72 | 385 ± 24 | 0.59; 0.59 |
| # GAD^67^ cells expressing OTR | 193 ± 37 | 148 ± 24 | 1.04; 0.36 |
| %GAD^67^ cells expressing OTR | 45 ± 5 | 38 ± 4 | 1.07; 0.35 |
| **Caudal DR** |  |  |  |
| # TPH cells | 93 ± 9 | 121 ± 39 | 0.71; 0.52 |
| # TPH cells expressing OTR | 23 ± 6 | 26 ± 7 | 0.28; 0.79 |
| %TPH cells expressing OTR | 25 ± 6 | 22 ± 3 | 0.52; 0.63 |
| # GAD^67^ cells | 201 ± 4 | 106 ± 16 | 5.67; 0.01* |
| # GAD^67^ cells expressing OTR | 34 ± 4 | 30 ± 5 | 0.63; 0.57 |
| % GAD^67^ cells expressing OTR | 17 ± 1 | 28 ± 4 | 2.87; <0.05* |

*Supplementary Table 1:* **Oxytocin receptor colocalization on serotonin, GABA, and dopamine neurons between diestrus virgins and recently-parturient dams.** Number of tryptophan hydroxylase (TPH), Glutamic acid decarboxylase^67^ (GAD^67)^, and tyrosine hydroxylase (TH) neurons expressing OTRs (Mean ± SEM) in female rats sacrificed as diestrus virgins or 3 hours after parturition. * indicates statistically significant group difference, p < 0.05.
