## Supplementary Table 2 for "Dorsal raphe oxytocin receptors regulate the neurobehavioral consequences of social touch"

| **Undisturbed Maternal Behavior**  **Tests** | **Scrambled (M ± SEM)** | **OTRKD (M ± SEM)** | **Group**  **(*F*_(1, 18)_*; p; η^2^_p)_*** | **Time**  **(*F*_(6)_*; p; η^2^_p)_*** | **Interaction**  **(*F*_6,108)_*; p; η^2^_p)_*** |
| --- | --- | --- | --- | --- | --- |
| Dam in nest | 1124 ± 24 | 916 ± 44 | 17.56; <0.01; 0.49 | 4.86; <0.01*; 0.21 | 2.57; >0.05; 0.13 |
| All Nursing | 918 ± 39 | 652 ± 56 | 15.16; <0.01; 0.46 | 0.82; 0.52; 0.04 | 3.91; 0.01; 0.18 |
| Kyphosis | 741 ± 37 | 455 ± 48 | 22.14; <0.01; 0.55 | 3.40; 0.01*; 0.16 | 3.54; 0.01; 0.17 |
| Supine nursing | 79 ± 20 | 76 ± 30 | 0.01; 0.92; <0.01 | 5.81; <0.01^;0.24 | 0.90; 0.45; 0.05 |
| Prone nursing | 97 ± 17 | 122 ± 12 | 1.33; 0.26; 0.07 | 1.72; 0.15; 0.09 | 1.36; 0.25; 0.07 |
| Hovering over the litter | 206 ± 31 | 264 ± 31 | 1.66; 0.21; 0.09 | 5.12; <0.01*; 0.16 | 1.01; 0.41; 0.05 |
| Erect postures (hovering over + kyphosis) | 947 ± 29 | 718 ± 40 | 21.19; <0.01; 0.54 | 8.68; <0.01*; 0.33 | 2.22 ; 0.07; 0.11 |
| Passive postures (supine + prone nursing | 177 ± 23 | 198 ± 35 | 0.25 ; 0.62; 0.01 | 3.42; 0.01^; 0.16 | 1.44; 0.23; 0.07 |
| Licking pups | 105 ± 19 | 140 ± 25 | 1.32; 0.27; 0.07 | 3.79; 0.01^; 0.17 | 0.81; 0.52; 0.04 |
| Retrieval | 0.9 ± 0.8 | 1.9 ± 1.1 | 0.52 ; 0.48; 0.03 | - | - |
| Non-pup directed behavior | 230 ± 26 | 442 ± 41 | 18.87; <0.01; 0.51 | 1.44; 0.22; 0.07 | 4.22; <0.01; 0.19 |
| Nesting | 13 ± 5 | 31 ± 8 | 3.29; 0.09; 0.16 | 1.66; 0.18; 0.09 | 1.54; 0.21; 0.08 |
| Nest quality | 38 ± 3 | 27 ± 5 | 4.67; 0.04; 0.21 | 4.55; <0.01*; 0.20 | 2.00; 0.10; 0.10 |
| Self-grooming | 72 ± 10 | 115 ± 12 | 7.80; 0.01; 0.30 | 0.84; 0.51; 0.05 | 4.04; <0.01; 0.18 |
| Ingestive behaviors | 29 ± 6 | 94 ± 14 | 19.22; <0.01; 0.52 | 2.02; 0.10; 0.10 | 2.27; 0.07; 0.11 |
| Exploring | 63 ± 10 | 136 ± 16 | 15.00; <0.01; 0.46 | 1.10; 0.36; 0.06 | 2.77; 0.04; 0.13 |
| Sleeping away from pups | 53 ± 12 | 56 ± 17 | 0.35; 0.56; 0.02 | 3.47; 0.01^; 0.16 | 0.56; 0.69; 0.03 |
| **Pup Retrieval Tests** |  |  |  |  |  |
| Latency to retrieve first pup (s) | 19 ± 5 | 27 ± 9 | 0.23; 0.83; 0.10 | - | - |
| Latency to group all pups (s) | 173 ± 23 | 186 ± 23 | 0.45; 0.66; 0.20 | - | - |
| Kyphosis | 0.6 ± 0.4 | 0.5 ± 0.3 | 0.31; 0.76; 0.14 | - | - |
| Hovering over litter | 16 ± 1 | 15 ± 1 | 1.14; 0.27; 0.53 | - | - |
| Licking pups | 9 ± 1 | 8 ± 1 | 1.52; 0.15; 0.67 | - | - |
| Non-pup directed behavior | 10 ± 1 | 11 ± 2 | 0.09; 0.93; 0.55 | - | - |
| Self-grooming | 3 ± 1 | 3 ± 0.4 | 0.94; 0.36; 0.40 | - | - |
| Exploring | 7 ± 1 | 8 ± 1 | 1.26; 0.22; 0.57 | - | - |

*Supplementary Table 2:* **Oxytocin receptor knockdown affected numerous maternal caregiving behaviors.** Frequencies (Mean ± SEM) of maternal behaviors displayed by scrambled and oxytocin receptor knockdown (OTRKD) dams during three 30-min observations each day on postpartum days 2 – 8 and following retrieval testing. ^ indicates increasing frequency of behavior across postpartum day. *indicates decreasing frequency of behavior across postpartum day.
